# Reduction of bundle sheath size boosts cyclic electron flow in C_4_ *Setaria viridis* acclimated to low light

**DOI:** 10.1101/2021.04.11.439306

**Authors:** Chandra Bellasio, Maria Ermakova

**Affiliations:** Department of Biology, University of the Balearic Islands, 07122 Palma, Illes Balears, Spain; Centre of Excellence for Translational Photosynthesis, Research School of Biology, The Australian National University, Acton, Australian Capital Territory 2601, Australia

**Keywords:** C_4_ photosynthesis, Kranz anatomy, light harvesting, bundle sheath, light reactions, carbon reactions, NADP-ME, gas-exchange, modelling

## Abstract

When C_4_ leaves are exposed to low light, CO_2_ concentration in the bundle sheath (BS) cells decreases, causing an increase in photorespiration relative to assimilation, and a consequent reduction in biochemical efficiency. These effects can be mitigated by complex acclimation syndromes, which are of primary importance for crop productivity, but not well studied. We unveil an acclimation strategy involving regulation of electron transport processes. Firstly, we characterise anatomy, gas-exchange and electron transport of C_4_ *Setaria viridis* grown under low light. Through a purposely developed biochemical model, we resolve the photon fluxes and reaction rates to explain how the concerted acclimation strategies sustain photosynthetic efficiency. Our results show that a smaller BS in low light-grown plants limited leakiness (the ratio of CO_2_ leak rate out of the BS over the rate of supply via C_4_ acid decarboxylation) but sacrificed light harvesting and ATP production. To counter ATP shortage and maintain high assimilation rates, plants facilitated light penetration through the mesophyll and upregulated cyclic electron flow in the BS. This shade tolerance mechanism based on optimisation of light reactions is potentially more efficient than the known mechanisms involving the rearrangement of carbon metabolism, and can potentially lead to innovative strategies for crop improvement.

**Significance:** We mechanistically link the optical cross-section of leaf compartments with the rate of electron transport, the engagement of cyclic electron flow, the relative rate of ATP and NADPH generation, and fluxes through the carbon metabolism. The striking capacity of *Setaria viridis* to counter the decrease in light absorption in the bundle sheath with an increase of cyclic electron flow presents perhaps the most efficient mechanism of shade acclimation.

## Introduction

Leaves of the majority of C_4_ plants are organised in concentric cylinders: an external tube of mesophyll (M) cells surrounds a tube of bundle sheath (BS) cells, which are wrapping the innermost vasculature (Figure 1). This spatial arrangement imposes that light and CO_2_ entering C_4_ leaves pass through M cells before reaching the BS. CO_2_ is hydrated to HCO_3_^-^ by carbonic anhydrase in the M cytosol and then fixed by PEP carboxylase (PEPC) into C_4_ acids, which may be transaminated or chemically reduced. C_4_ acids diffuse to BS cells where they are decarboxylated either primarily by NADP-malic enzyme (NADP-ME), NAD-malic enzyme (NAD-ME), PEP carboxykinase (PEPCK), or by a combination of these. Decarboxylation provides higher CO_2_ partial pressure at the site of ribulose 1,5-bisphosphate carboxylase/oxygenase (Rubisco) thereby largely suppressing the competitive reaction with oxygen leading to the wasteful process of photorespiration (von Caemmerer and Furbank, 2003, Bellasio *et al.*, 2014). For this function, C_4_ photosynthesis is also referred to as biochemical carbon concentrating mechanism (CCM). Pyruvate resulting from the C_4_ acids decarboxylation diffuses back to M cells where it is regenerated into PEP by pyruvate orthophosphate dikinase (PPDK) using 2 ATP molecules. Some of the CO_2_ released in BS cells diffuses out to M cells (called leakage) which incurs in an additional ATP cost of C_4_ photosynthesis (Farquhar, 1983, Henderson *et al.*, 1992, Bellasio and Griffiths, 2014a).

**Figure 1.**
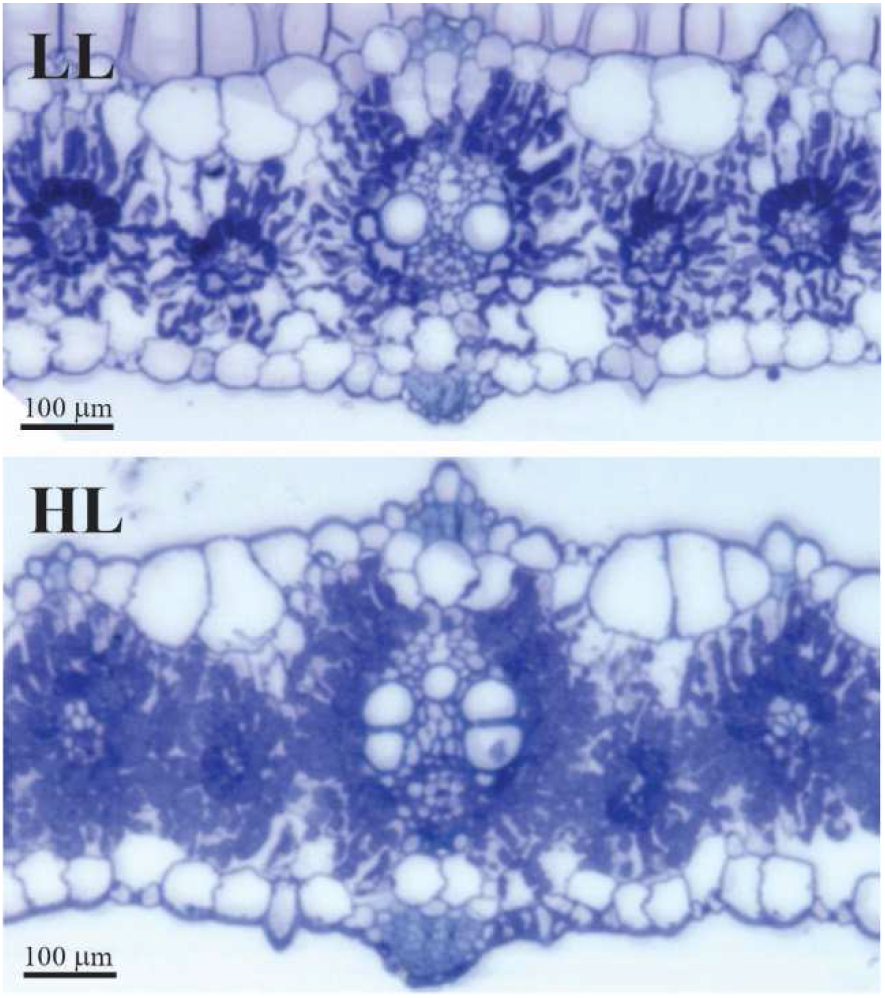
Light microscopy images of the leaf cross-sections used for anatomical measurements from *S. viridis* grown under high light (1000 μmol m^-2^ s^-1^, HL) or low light (300 μmol m^-2^ s^-1^, LL).

C_4_ photosynthesis is highly sensitive to limiting light intensities. Under low irradiance, ATP and NADPH generation slows down, limiting both the reductive pentose phosphate cycle (RPP) and the C_4_ cycle. Since activities of RPP and the C_4_ cycle are tightly balanced (Kromdijk *et al.*, 2014) through light-regulation (Bailey *et al.*, 2007), a slowdown of C_4_ acids diffusion into the BS inevitably results in a decrease of CO_2_ concentration in BS cells (*C*_BS_). Consequently, the ratio between CO_2_ and O_2_ concentration at the active sites of Rubisco decreases, favouring oxygenation over carboxylation, lowering the biochemical efficiency [ATP per gross assimilation of CO_2_] of C_4_ photosynthesis (Furbank *et al.*, 1990, Ubierna *et al.*, 2013)]. In crop canopies, where up to 50 % of net CO_2_ uptake is fixed by shaded leaves, (Baker *et al.*, 1988, Long, 1993) light limitation plays an important role in decreasing canopy productivity (Kromdijk *et al.*, 2010), and understanding acclimation strategies of C_4_ metabolism to light limitation is critical to increase crop production to meet the raising food and fodder demand (Evans *et al.*, 1991, Bellasio and Griffiths, 2014a).

Shading limits C_4_ photosynthesis also at the cell level. M and BS cells both need light to power photosynthesis. The amount of light that can be harvested by the BS cell is limited by their size [precisely by the BS optical cross section (Bellasio and Lundgren, 2016)], by light absorption in the M [because of the concentric organisation, BS cells are in effect shaded, meaning that light harvesting in the M reduces the light available to reach BS cells (Kramer and Evans, 2011)], and depends on light quality. Green light is not strongly absorbed by chlorophyll, meaning that it can easily reach the BS, while blue light is all taken up by the upper chloroplasts and preferentially excites M cells (Evans *et al.*, 2007). M cells use light to produce ATP and NADPH through linear electron flow (LEF) engaging both photosystems (PS) while BS cells of NADP-ME plants are thought to produce mainly ATP through cyclic electron flow (CEF) around PSI. In this way, the relative absorption of excitation energy will influence the availability of ATP and NADPH. Bellasio and Griffiths (2014c) proposed that these environmentally driven shifts in ATP and NADPH production can be countered by adjusting ATP and NADPH consumption by carbon metabolism, with a surprising degree of flexibility – but only up to a defined limit, ultimately determining how the biochemical work is apportioned between the two cell types. However, the possibility that plants could respond to shift in energy availability by rearranging light reactions, in such a way that also the production of ATP and NADPH is regulated, received little attention.

We have recently shown that the photosynthetic apparatus of *Setaria viridis*, a model C_4_ plant of NADP-ME subtype, rearranges in response to low light in a cell-specific fashion, preferentially increasing electron transport capacity in BS cells (Ermakova *et al.*, 2021b). We hypothesized that this was a response to a shortage of ATP in the BS, which could depend either on an increased consumption from dark reactions, or on a lower rate of production caused by a shortage of light. Here we resolve these drivers and their causal relationship. First, we measure anatomical properties of leaves from *S. viridis* grown under high light (HL) and low light (LL) and estimate how these would influence light harvesting of the M and the BS. Then, we conduct a comprehensive gas exchange characterisation under ambient and low O_2_, with concurrent determination of PSII and PSI effective quantum yields, to estimate biochemical parameters and conductance to CO_2_ diffusion at the M/BS interface (*g*_BS_). Finally, we develop a model integrating two electron transport chains with the light-harvesting properties of M and BS to study how the apportioning of light and carbon metabolism shifts in LL plants, and how these changes affect biochemical efficiency. We demonstrate that the decreased light interception in BS cells is effectively counteracted by an increase of CEF, uncovering another important link between biochemical properties and structural characteristics of C_4_ leaves.

## Methods

### Plants

*S. viridis* plants (A10 ecotype) were grown in 2 L pots filled with commercial soil mix (Debco, Tyabb, Australia) supplemented with 1 g L^-1^ of slow release fertilizer (Osmocote, Scotts, Bella Vista, Australia). Plants were grown in controlled chambers with 28°C day, 24°C night, 60 % humidity, and 16 h illumination at the intensity of 1000 μmol m^-2^ s^-1^ or 300 μmol m^-2^ s^-1^ provided by halogen incandescent lamps (42 W, 2800 K, warm white, clear glass, 630 lumens, CLA, Brookvale, Australia) and Pentron Hg 4 ft fluorescent tubes (54 W, 4100 K, cool white, Sylvania, Wilmington, MA, USA), and reached by hanging a shade cloth above some of the plants. Growth light spectra are shown in Figure S2. All measurements were performed on the youngest fully expanded leaves sampled before flowering between 15 and 25 days after germination.

### Leaf anatomy

Resin–embedded cross–sections were prepared and imaged according to Pengelly *et al.* (2010). Quantification of anatomical parameters was performed using Image J software (NIH, WI) on an equal number of secondary and tertiary veins as described in Bellasio and Lundgren (2016). Calculations of the M surface area exposed to the intercellular airspace (S_m_) and the BS surface area per unit of leaf area (S_b_) were made as described in Pengelly *et al.* (2010) using the curvature correction factor of 1.43 from Evans *et al.* (1994). Apparent absorbance was calculated as the log10 of the ratio between the luminance of background divided by the luminance of the tissue, both processed with Image J as described in Bellasio and Lundgren (2016) and averaged over two regions for *n*=6 replicates. Leaf absorptance was the complement to one of leaf reflectance and transmittance, both measured with a Li-Cor 1800-12 (Li-Cor, Lincoln, NE) integrating sphere, coupled to a Li–190S light sensor, following the manufacturer’s instructions for calibration and calculations.

### Chlorophyll

Total chlorophyll was extracted from frozen leaf discs ground using TissueLyser II (Qiagen, Venlo, The Netherlands) in 80% acetone, buffered with 25 mM 4-(2-hydroxyethyl)-1-piperazineethanesulfonic acid (Hepes)-KOH (pH 7.8). Chlorophyll *a* and *b* content was measured at 750 nm, 663.3 nm and 646.6 nm, and calculated according to Porra *et al.* (1989). The fraction of total leaf chlorophyll in BS cells was determined from the Chlorophyll *a*/*b* ratios leaves, BS and M cells as described in Ermakova *et al.* (2021b).

### Western blotting and immunolocalization

Protein isolation from leaves, gel electrophoresis and western blotting were performed as described in Ermakova *et al.* (2019). PEPCK antibodies (Agrisera, Vännäs, Sweden) were used according to the manufacturer’s protocol.

Immunodetection of PPDK on lightly fixed leaf cross-section was performed according to (Ermakova *et al.*, 2021a). Sections were treated with 1:100 PPDK primary antibody (Karki *et al.*, 2020), 1:200 Alexa Fluor 488–conjugated goat anti-rabbit secondary antibody (Life Technologies, Eugene, OR) and 0.05 % calcofluor white to stain cell walls. Fluorescence signal was captured with the Leica DM5500 microscope (Leica, Wetzlar, Germany) equipped with the Leica DFC7000T camera using the Leica Application Suite 4.12 software. Fluorescence was detected at 505–565 nm for PPDK (excitation 490–510 nm, YFP Filter Cube, Leica) and at 450–490 nm for cell walls (excitation 330–370 nm, A4 Filter Cube, Leica).

### Simultaneous gas exchange, chlorophyll fluorescence and P700 measurements

Gas exchange, fluorescence and P700^+^ absorbance changes were measured simultaneously with the setup of Bellasio and Farquhar (2019) on *n* = 3 biological replicates. Briefly, a portable gas exchange system (LI6400XT, Li–Cor) was modified to operate at low CO_2_ concentrations (see licor.com) and fitted with a 6400–06 PAM2000 adapter (courtesy of Susanne von Caemmerer), holding a fibre probe in the upper leaf cuvette distant enough to minimise shading. Leaves were aligned without overlapping their edges to fill the cuvette. Light was provided by a bespoke red–blue light source, regulated to provide circa 90% red and 10% blue light, positioned to illuminate the leaf uniformly. Light intensity was measured through an in–chamber Gallium arsenide photodiode, calibrated using a Li–250 light sensor (Li–Cor). Neoprene gaskets were used on both sides of the cuvette. A mixture of 2 % O_2_ was prepared by mixing ambient air and N2 with a bespoke gas mixing unit (kindly assembled by Suan Chin Wong). This mix of ambient air was CO_2_-scrubbed with soda lime and humidified to a dew point of 15–17 °C upstream of the inlet to maintain water vapour pressure deficit around 1 kPa. CO_2_ was added from a large cylinder (BOC, St Peters, Australia), using the CO_2_ injection unit of the LI6400XT. Mass flow leaks (Boesgaard *et al.*, 2013) were monitored with a gas flow meter as detailed in Bellasio *et al.* (2016b).

PSII and PSI yields were measured with a Dual PAM–F (Heinz Walz GmbH, Effeltrich, Germany, courtesy of Dean Price). PSII activity was assessed with the pulse amplitude modulated fluorescence method using 620 nm measuring light of 9 μmol m^-2^ s^-1^ (Schreiber *et al.*, 1986). The redox state of P700, the reaction centre of PSI, was assessed by detecting absorbance of the cation at 830 nm with a dual wavelength (830/875 nm) unit, following the method of Klughammer and Schreiber (1994). Saturating pulses of red light (635 nm) at 20,000 μmol m^-2^ s^-1^ were used. First, P_M_, the maximal level of P700^+^, was recorded upon the application of a saturating pulse on top of the far-red light, and P0, the minimal P700^+^ signal, was recorder after that saturating pulse. Photosynthetic parameters (P, the steady-state P700^+^ signal; P_M′_, the maximal P700^+^ signal under light; F, the steady-state fluorescence signal; F_M′_, the maximal fluorescence signal under light) were monitored by the saturating pulse application at different irradiances, CO_2_ and O_2_ concentrations. The effective quantum yield of PSII [*Y*(*II*)] was calculated according to Genty *et al.* (1989). The effective quantum yield of PSI [*Y*(*I*)] and the non-photochemical yields of PSI caused by donor [*Y*(*ND*)] or acceptor [*Y*(*NA*)] side limitation were calculated as described in Klughammer and Schreiber (2008).

Four photosynthetic response curves (*A/PPFD* and *A*/*C*_i_ curves, under ambient and low [O_2_]) were measured at 25 °C in daylong experiments. *A*/*C*_i_ curves were measured first, under a *PPFD* of 1000 μmol m^-2^ s^-1^ and imposing reference CO_2_ concentrations of 20, 40, 60, 80, 100, 200, 300, 400, 500, 600, 800, 1000 μmol mol^-1^ and a minimum of 120 s between steps; *A*/*PPFD* curves were measured under reference [CO_2_] of 420 μmol mol^-1^ imposing *PPFD*s of 1500, 1000, 200, 500, 150, 100, 75, 50, 30 μmol m^-2^ s^-1^ and a minimum of 180 s between steps. The order between ambient and low O_2_ was inverted each day. Flow rate was 400 μmol s^-1^; CO_2_ diffusion through the gaskets was compensated by lengthening the tubing of the LI6400XT reference gas (Bellasio and Farquhar, 2019). Gas response curves were analysed using the protocol of Bellasio *et al.* (2016a). CO_2_ concentration at the M carboxylation sites was calculated as *C*_M_ = *C*_i_ - *A*/*g*_M_, where mesophyll conductance (*g*_M_) was 1.5 for HL and 1.2 mol m^-2^ s^-1^ for LL derived by adjusting for Sm (Table 1) the data of Ubierna *et al.* (2017).

**Table 1.**
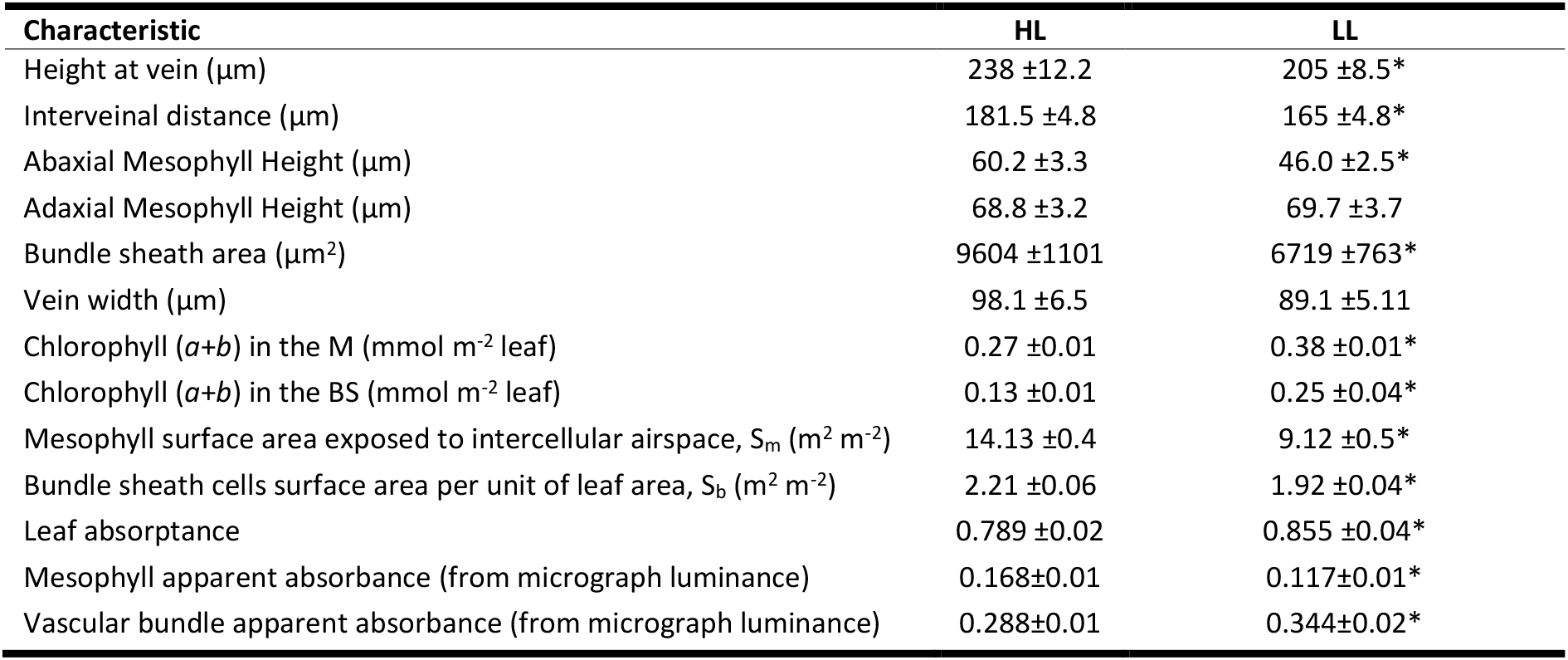
Anatomical and biochemical characteristics of leaves, mesophyll (M) and bundle sheath (BS) cells of *S. viridis* grown at high light (1000 μmol m^-2^ s^-1^, HL) or low light (300 μmol m^-2^ s^-1^, LL). Mean values are shown ± SE, *n* = 20 (except for chlorophyll and leaf absorptance *n* = 3, apparent absorbance *n* = 6). Asterisks indicate statistically significant difference between two light regimes (*P* < 0.05).

| Characteristic | HL | LL |
| --- | --- | --- |
| Height at vein ($\mu\text{m}$) | 238 ±12.2 | 205 ±8.5* |
| Interveinal distance ($\mu\text{m}$) | 181.5 ±4.8 | 165 ±4.8* |
| Abaxial Mesophyll Height ($\mu\text{m}$) | 60.2 ±3.3 | 46.0 ±2.5* |
| Adaxial Mesophyll Height ($\mu\text{m}$) | 68.8 ±3.2 | 69.7 ±3.7 |
| Bundle sheath area ($\mu\text{m}^2$) | 9604 ±1101 | 6719 ±763* |
| Vein width ($\mu\text{m}$) | 98.1 ±6.5 | 89.1 ±5.11 |
| Chlorophyll ($a+b$) in the M ($\text{mmol m}^{-2}$ leaf) | 0.27 ±0.01 | 0.38 ±0.01* |
| Chlorophyll ($a+b$) in the BS ($\text{mmol m}^{-2}$ leaf) | 0.13 ±0.01 | 0.25 ±0.04* |
| Mesophyll surface area exposed to intercellular airspace, $S_m$ ($\text{m}^2 \text{m}^{-2}$) | 14.13 ±0.4 | 9.12 ±0.5* |
| Bundle sheath cells surface area per unit of leaf area, $S_b$ ($\text{m}^2 \text{m}^{-2}$) | 2.21 ±0.06 | 1.92 ±0.04* |
| Leaf absorptance | 0.789 ±0.02 | 0.855 ±0.04* |
| Mesophyll apparent absorbance (from micrograph luminance) | 0.168±0.01 | 0.117±0.01* |
| Vascular bundle apparent absorbance (from micrograph luminance) | 0.288±0.01 | 0.344±0.02* |

### C_4_ photosynthesis model

Biochemical models may include formulations describing a variety of processes (*e.g.*, electron transport), but, at any given time, only the formulation describing the process that is limiting photosynthesis in that circumstance will be representative (outputs are called ‘actual’ rates), while others are not [outputs are called ‘potential’ rates (Farquhar *et al.*, 1980)]. The mismatch between formulations can sometimes be useful, for instance, to estimate stomatal conductance in C_3_ (Farquhar and Wong, 1984) and C_4_ plants (Bellasio *et al.*, 2017), but typically it generates ambiguity in curve fitting [*e.g.*, fluorescence, reviewed in Bellasio *et al.* (2016b)] and difficulties in locating the cut-off points (Gu *et al.*, 2010). To avoid discontinuities, we merged enzyme limitations into a light-limited model by introducing a function describing the quenching of *Y*(*II*) depending on *C*_M_, analogous to the classical *PPFD*-dependence of electron transport rate. This function depends on the familiar biochemical quantities *V*_PMAX_ and *K*_P_, but we have not derived an explicit expression at this stage.

The model is organized to simulate two compartments representing M and BS cells (Figure 2): each harvests light to drive a distinct electron transport chain (Note S1, Figure S1). In both electron transport chains, the ratio between the rates of ATP and NADPH production is varied via adjustment of CEF (through the parameter *f*_Cyc_), which may flow through PGR5/PGRL1 or the NAD(P)H dehydrogenase-like complex (NDH, adjusted through the parameter *f*_NDH_). When *f*_Cyc_ is adjusted, the proportion of light absorbed by photosystems varies to maintain invariant the sum of light absorbed by the two [equations described in supporting information of Yin *et al.* (2004)]. This is of key importance when modelling energetics and CEF engagement under limiting light (Bellasio, 2019) because, otherwise, supplemental CEF would be driven by an increase in light absorbed [but *s* would be invariant, main text of Yin *et al.* (2004)]. The reducing power requirements for nitrogen reduction and for the water-water cycle are explicitly accounted for as a fraction of pseudo-cyclic electron flow, after Yin and Struik (2012), but we did not use the functionality in this study. A key innovation is that the rate of oxygen evolution in the BS is calculated from the actual rate of electron transport in both compartments, instead of being assumed to scale to assimilation through the parameter α as in all the models derived from Berry and Farquhar (1978).

**Figure 2.**
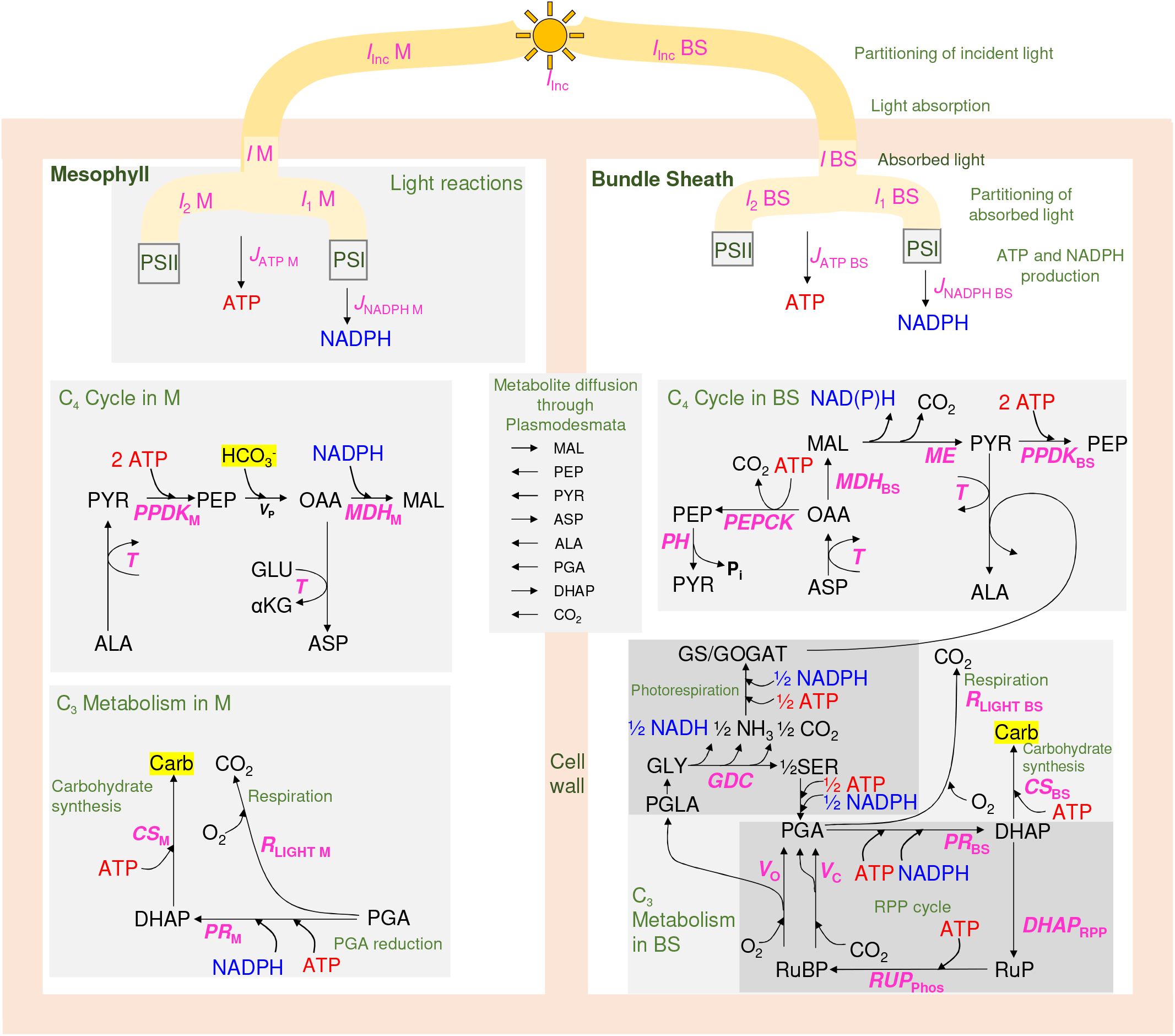
Schematic of the processes included in the model. Metabolites are in black; fluxes are schematised by black arrows and pink symbols which directly link to equations reported in Supporting Notes; key processes are briefly described in bright green. The leaf is divided into M and BS compartments. Incident *PPFD* (*I*_inc_) may reach M (*I*_Inc M_) or BS cells (*I*_Inc BS_) depending on anatomical characteristics and size of the light-harvesting machinery. A fraction *I* is absorbed by PSI or PSII (*I*_1_ or *I*_2_, respectively). Light reactions (for more details see Figure S1) result in the production of NADPH and ATP, which are consumed by carbon metabolism encompassing C_4_ and C_3_ activity. The C_4_ cycle reactions appear in the centre of the M and BS compartments. CO_2_ is initially hydrated to bicarbonate (the point of entry is highlighted in yellow) and fixed by PEP carboxylase (PEPC) at the rate *V*_P_ to form oxaloacetate (OAA). This may be reduced to malate (MAL) or to aspartate (ASP) through transamination (*T*), *i.e.*, the exchange of amino-groups with glutamate (GLU). MAL and ASP both diffuse to BS cells; MAL is decarboxylated with the concurrent reduction of NADP by malic enzyme (ME) while ASP is deaminated to OAA (the exchange of amino-groups with alphaketoglutarate (αKG) is implied) and may be decarboxylated by PEP carboxykinase (PEPCK) or by malic enzyme (ME). The regeneration of PEP is shared between M and BS cells depending on energy availability. The C_3_ metabolism appears at the bottom, partitioned between M and BS compartments. Rubisco carboxylation and oxygenation reactions (*V*_C_ and *V*_O_) consume RuBP and produce PGA and PGLA and are fully compartmentalised to BS cells. PGLA is recycled through the photorespiration cycle eventually regenerating PGA. This is consumed by respiration (*R*_LIGHT_), assumed to be entirely supplied by newly assimilated PGA, and is reduced (*PR*) to triose phosphate (DHAP), which is a substrate of carbohydrate synthesis (CS). The final product of photosynthesis is a generic triose carbohydrate (highlighted in yellow). The majority of DHAP enters the sugar conversion phase of the reductive pentose phosphate (RPP) cycle, exclusive to the BS. Metabolites for which fluxes are calculated are listed in the middle, and assumed to be positive when occurring in the ordinary direction indicated by arrows. The Excel workbook provided renders outputs according to this scheme.

ATP and NADPH drive the reactions of the carbon metabolism, described following Bellasio and Griffiths (2014c), McQualter *et al.* (2016), and Bellasio (2017). While the NADPH available in the BS is consumed by the RPP and the photorespiratory cycles, the ATP available in the M and any ATP surplus in the BS are consumed by the C_4_ cycle, until all the ATP and NADPH available have been consumed. In this way, the partitioning between cycles and the apportioning of metabolic work between M and BS cells emerges directly from the rate of ATP and NADPH produced by light reactions, instead of being assumed through the parameter *x* as in all the approaches following Berry and Farquhar (1978). Differently from Bellasio (2017), where all intermediate photosynthetic types were modelled, here C_4_ photosynthesis is fully expressed, such that the glycine decarboxylase complex (GDC) and Rubisco are both fully compartmentalised to BS cells (Bellasio, 2017). However, C_4_ subtypes can be changed seamlessly, forming a continuous biochemical space that can be explored by varying two parameters: *f*_MDH_, representing the engagement of MDH in M cells and controlling the trajectory between the NADP-ME subtype and the NAD-ME subtype, and *f*PEPCK, representing the capacity of PEPCK to consume ATP available in BS cells and representing the level of PEPCK expression relative to other carboxylases (we will explore these functionalities in dedicated studies). In the decarboxylation of malate, the co-products NADH and NADPH are assumed to be equivalent or interconvertible. This may be underpinned by an actual biochemical interconversion, mediated by the intercellular operation of the malate shuttle (Furbank, 2011), or simply by the capacity of malic enzyme (ME) to dock both cofactors, as suggested by early studies (Kanai and Edwards, 1973).

M and BS cells are connected by an exchange of metabolites (malate, MAL; phosphoenolpyruvate, PEP; pyruvate, PYR; aspartate, ASP; alanine, ALA; 3-phosphoglyceric acid, PGA; dihydroxyacetone phosphate, DHAP; and CO_2_) which diffuse obeying mass balance constraints. Diffusion is non-limited, except that of CO_2_, restrained by a finite conductance (*g*_BS_) that allows higher CO_2_ concentration in BS cells (*C*_BS_) based on classical underpinnings (Berry and Farquhar, 1978). Glutamate and αKG do not diffuse directly between M and BS cells, but exchange amino-groups with ALA and PYR (Pick *et al.*, 2011, Mallmann *et al.*, 2014, Schlüter *et al.*, 2019). If reactions can proceed in opposite direction, they are assumed to proceed only in the direction with positive balance. Respiration is a shared process between M and BS cells. To avoid a futile cycle consisting of concurrent glycolysis and carbohydrate synthesis, respiration is assumed to be supplied by new assimilates (PGA) in line with Bellasio and Griffiths (2014c). The ATP and NADH produced during respiration are neglected, assumed to be consumed by basal metabolism. Any ATP and NADH residual imbalances produced during respiration in the light are absent, supported by the analysis of Buckley and Adams (2011) who proposed they would be dissipated by alternative oxidases. The final product of photosynthesis is a generic triose carbohydrate, whose destiny is not further followed by the model.

Details of the functions describing reaction rates and CO_2_ and O_2_ concentrations in M and BS compartments are provided in Supporting Note S2; reducing power poise is in Supporting Note S3 and metabolite flows in Note S4. Model sensitivity analysis is shown in Table S2.

### Parameterisation

Parameters were either derived from measurements (in which case data was randomized and averaged for HL and LL plants), taken from dedicated studies, or assumed. Light penetration in the leaf was simulated through a separate optical model, freely downloadable from Supporting Information of Bellasio and Lundgren (2016). The number of layers of adaxial M cells was 289 for HL plants and 340 for LL plants; the number of layers of the vein was 458 for HL plants and 435 for LL plants; the number of layers of abaxial M cells was 253 for HL plants and 225 for LL plants; the fraction of interveinal distances occupied by the vein was 0.56 for HL plants and 0.532 for LL plants, all derived from anatomical measurements (Table 1). The absorbance of the adaxial M cells was set to equal that of M cells following the observation that BS extensions are absent in *S. viridis;* the absorbance of the vein relative to M cells 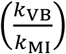 was 1.65 for HL plants and 2.73 for LL plants, calculated by dividing the measured BS/M ratio of chlorophyll content per metre square of leaf, by the measured ratios of BS/M areas, 0.29 for HL and 0.24 for LL; reflectance was 0.067 for HL plants and 0.081 for LL plants (Table 1).

The combined model of light and dark reactions has 29 input quantities. Three define the environmental characteristics (*PPFD*, *C*_M_ and *O*_M_), 17 define light reactions (*Y*(*II*)_0_, *s*, *f*_Pseudocyc NR M_, *f*_Pseudocyc NR BS_, *f*_Q_, *h*, *Y*(*I*)_LL_ *f*_NDH M_, *f*_NDH BS_, *f*_Cyc M_, *f*_Cyc BS_, α_C_, *V*_0 C_, θ_C_, α_v_, *V*_0 V_, θ_v_), BS six define dark reactions *f*_PH_, *f*_PEPCK_, *f*_RLIGHT_, *f*_CS_, *f*_MDH_), and three more (*g*_BS_, 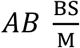, *R*_LIGHT_) are not grouped. 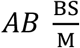 was taken from the light penetration model described above, *R*_LIGHT_ and *g*_BS_ were derived from gas exchange and fluorescence measurements with the procedure described in (Bellasio *et al.*, 2016a) (Table S1). *Y*(*II*)_0_ was derived by slightly increasing maximum values of *Y*(*II*)_LL_ so that Eqn S11 would return *Y*(*II*)_LL_ under ambient CO_2_ concentration and zero *PPFD.* α_C_, *V*_0 C_, θ_C_, α_V_, *V*_0 V_, θ_V_ were derived by fitting fluorescence data shown in Figure 3. *s*_0_ is the theoretical maximum value of *s* when *f*_Cyc_ is zero was adjusted for modelled 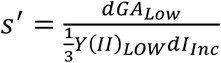 to fit measured values of *s*′ (Table S1). The *Q*–cycle was assumed obligate making *f*_Q_=1; *Y*(*I*)_LL_ was assumed 1 after Yin and Struik (2012). The behaviour of *f*_Pseudocyc NR BS_ and *f*_Pseudocyc NR M_ in response to CO_2_ and *PPFD* is not well known, therefore we assumed a low value for simplicity. The stoichiometry of the ATP synthase was set at *h*=4.67 H^+^ per ATP after structural evidence (Vollmar *et al.*, 2009, Hahn *et al.*, 2018). The operation of NDH is essential for C_4_ photosynthesis, but estimates for the electron flow through the complex are unknown, and difficult to imagine. Here *f*_NDH_ was set to 0.4 in M cells and 0.7 in BS cells following the observation that NDH- and PGR5-mediated electron flows are both operating in C_4_ plants (Takabayashi et al. 2005) while keeping in mind that the relative ratio of NDH to PSI was higher in BS cells than in M cells (Ermakova *et al.*, 2021b). *f*_Cyc M_ and *f*_Cyc BS_ were fitted to maximise *A* at each light intensity subject to the constraint that the activity of PPDK in BS cells is zero; fitted values are shown in Figure S3. *f*_RLIGHT_ and *f*_CS_ were set 0.5 after von Caemmerer (2000) and Bellasio (2017). *f*_PEPCK_ was set to zero to match immunoblotting (Figure 5). *f*_PH_ is irrelevant under the previous assumption. *f*_MDH_ sets the transition between NAD–ME and NADP–ME subtypes and was assumed to be 1 (Gutierrez *et al.*, 1974, John *et al.*, 2014).

**Figure 3.**
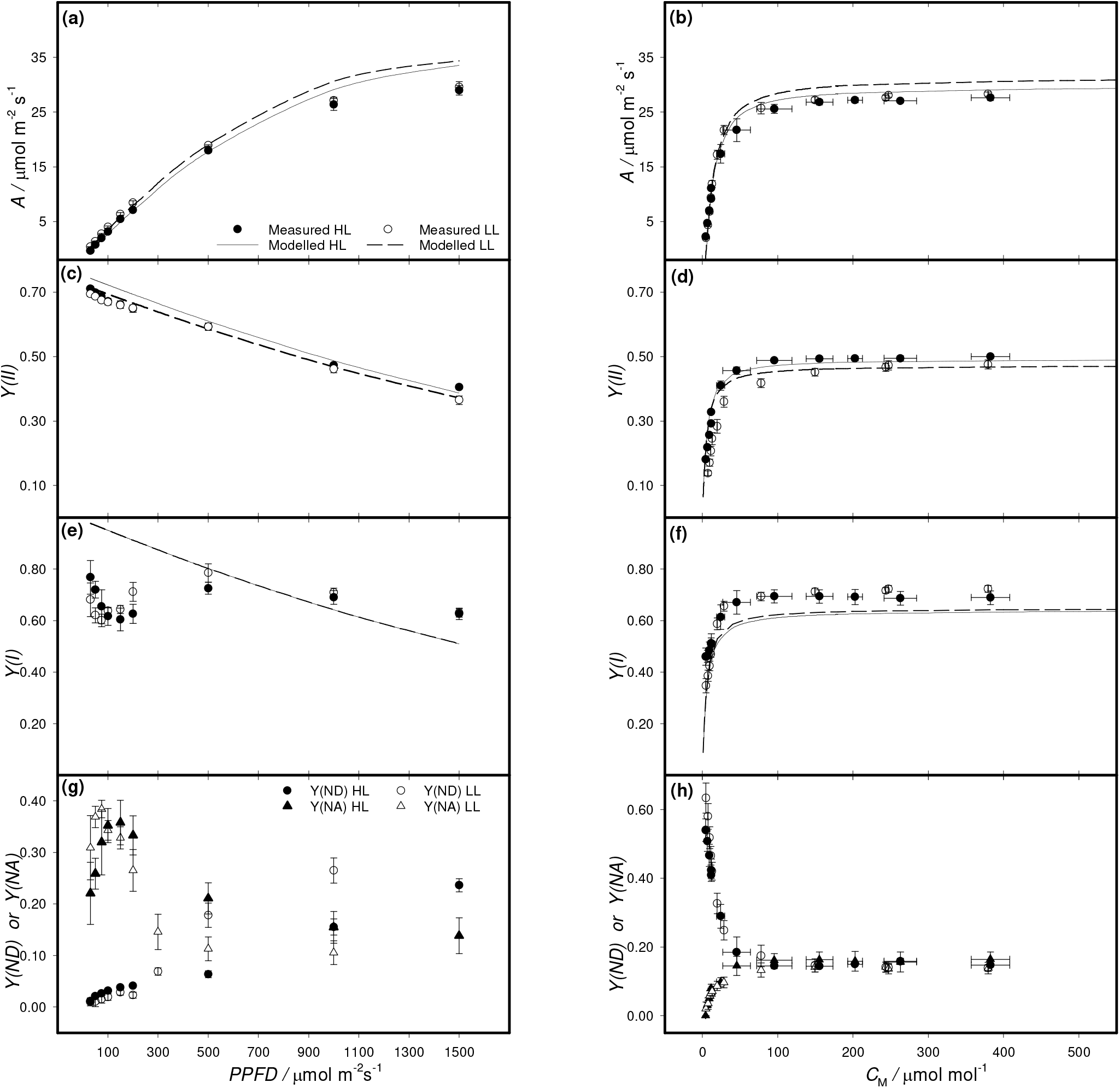
Measured and modelled gas exchange and photosystems yield under ambient O_2_ level. Symbols show response curves of CO_2_ assimilation (**a**, **b**), quantum yield of Photosystem II (**c**, **d**), quantum yield of Photosystem I (**e**, **f**), Photosystem I donor [*Y*(*ND*)] and acceptor side [*Y*(*NA*)] limitations (**g**, **h**) obtained for *S. viridis* grown at high light (HL, solid circles) and low light (LL, empty circles). Light curves measured under a reference [CO_2_] of 420 μmol mol^-1^ are shown in left panels, and CO_2_ response curves obtained under constant irradiance of 1000 μmol m^-2^ s^-1^ are shown in right panels. Mean ± SE, *n* = 3 biological replicates. *C*_M_, CO_2_ concentration in M cells. Corresponding curves obtained by switching the background gas to low O_2_ are shown in Figure 4. Lines show modelled responses obtained through a combined biochemical model of light reactions and carbon metabolism of C_4_ photosynthesis. Model parameters are listed in Table 2.

### Statistical analysis

The relationship between mean values for HL and LL plants was tested using a two-tailed, heteroscedastic Student’s *t*-test at *P* < 0.05 (Microsoft Excel® 2016). Error propagation in the model was not analysed.

## Results

### Leaf anatomy and chlorophyll

In HL plants, BS chloroplasts were uniformly distributed inside the cells (Figure 1). In LL plants, BS chloroplasts had clear centrifugal position whilst M chloroplasts were specifically arranged along the cell walls, particularly in adaxial M cells. This was captured by a higher apparent absorbance of M cells of HL plants (Table 1). Increased number of chloroplasts in adaxial BS cells compared to abaxial ones was more prominent in LL plants. In line with the higher chlorophyll content in the BS of LL plants relative to the M, LL vascular bundles showed increased apparent absorbance (Table 1). LL leaves had 15 % lower M thickness as well as 10 % lower interveinal distance compared to HL plants. The BS surface area per leaf area (S_b_) was 13 % lower, and the M surface area exposed to the intercellular airspace (S_m_) decreased by 35 %, in LL plants compared in HL plants.

### Gas-exchange and yield of photosystems

When the light-response was measured at 420 μmol mol^-1^ CO_2_ and 21 % O_2_, no difference was significant between HL and LL plants in CO_2_ assimilation rate (*A*) or effective quantum yields of PSII [*Y*(*II*)] and PSI [*Y*(*I*)] (Figure 3). Yields of non-photochemical losses in PSI due to the donor [*Y*(*ND*)] and acceptor [*Y*(*NA*)] side limitations did not differ between the two growth regimes (Figure 3g). At 2 % O_2_, no differences in light responses between HL and LL plants were detected (Figure 4).

**Figure 4.**
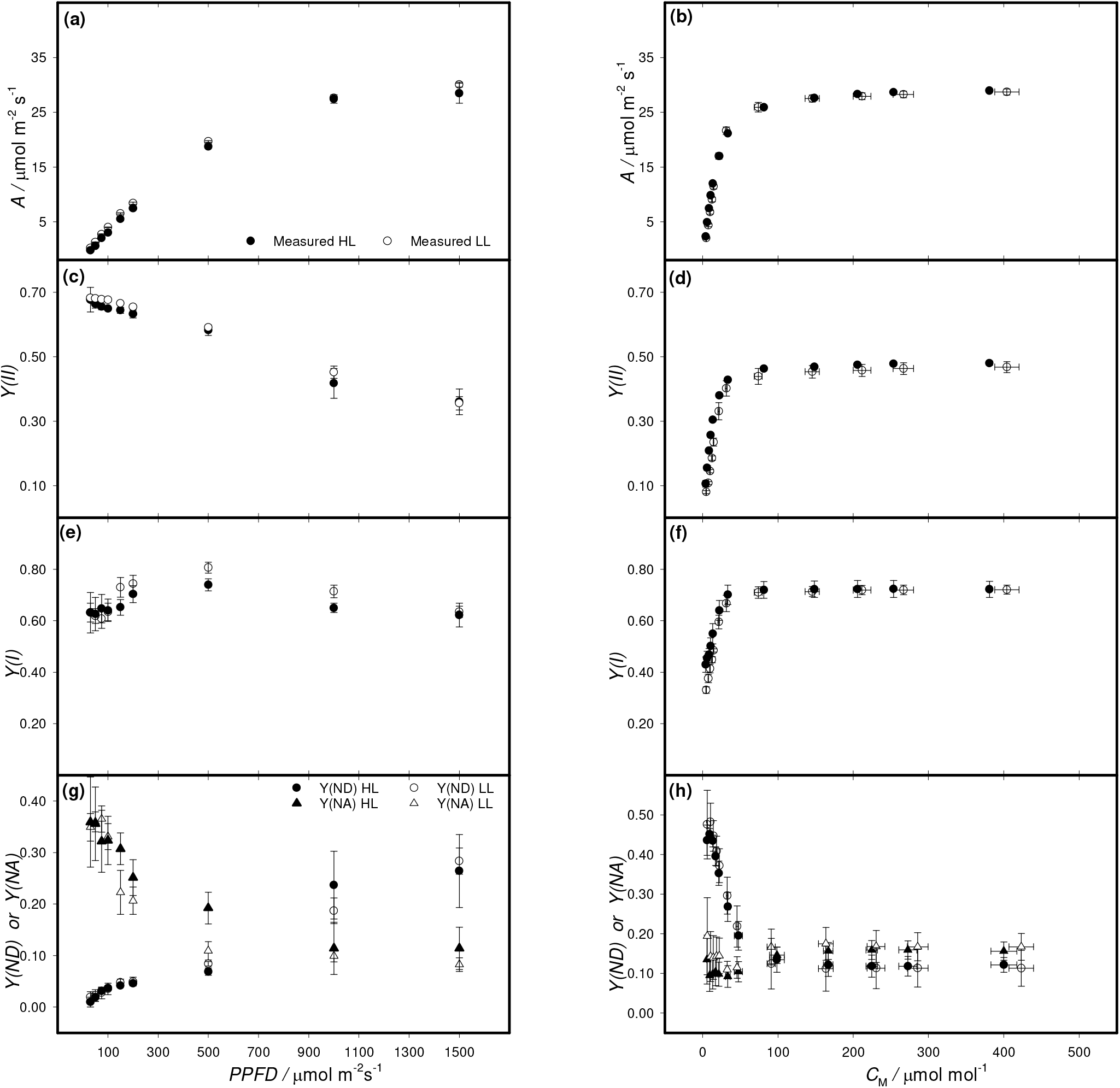
Measured gas exchange and photosystems yield at 2 % O_2_. Light curves measured under a reference [CO_2_] of 420 μmol mol^-1^ are shown in left panels, CO_2_ response curves obtained under constant irradiance of 1000 μmol m^-2^ s^-1^ are shown in right panels. Symbols show response curves of CO_2_ assimilation (**a**, **b**), quantum yield of Photosystem II (**c**, **d**), quantum yield of Photosystem I (**e**, **f**), Photosystem I donor side [*Y*(*ND*)] and acceptor side [*Y*(*NA*)] limitations (**g**, **h**) obtained for *S. viridis* grown at high light (HL, solid circles) and low light (LL, empty circles). Mean ± SE, *n* = 3 biological replicates. *C*_M_, CO_2_ concentration in M cells.

CO_2_ response curves were measured at a *PPFD* of 1000 μmol m^-2^ s^-1^ either at 21 % or 2 % O_2_. No differences in CO_2_ assimilation, *Y*(*II*), *Y*(*I*), *Y*(*ND*) or *Y*(*NA*) were significant between HL and LL plants independently of O_2_ level (Figure 3, Figure 4). Both HL and LL plants showed decreased *Y*(*II*) and increased *Y*(*ND*) at low *C*_M_, likely, representing an induction of non–photochemical quenching in the PSII antennae in response to the acidification of the thylakoid lumen. The latter is induced by the downregulation of ATP synthase activity in response to the reduced ATP consumption by the RPP cycle (Kanazawa and Kramer, 2002). Interestingly, *Y*(*NA*) at low *C*_M_ was higher when measured at 2 % O_2_ (Figure 3h, Figure 4h), pointing to a contribution of O_2_ to oxidating PSI via photorespiration and/or O_2_ photoreduction (Sagun *et al.*, 2021).

To aid interpretation of light and CO_2_ response curves, we fitted empirical and mechanistic models after Bellasio *et al.* (2016a). Fitted parameters in Table S1 show that, under ambient O_2_, the main differences were dependent on a decrease of respiration in the light (*R*_LIGHT_), light compensation point (*LCP*), CO_2_ compensation point (Γ) and BS conductance to CO_2_ diffusion (*g*_BS_) in LL plants. Curiously, O_2_ sensitivity of *R*_LIGHT_, quantum yield [*Y*(*CO*_2_)_LL_] and carboxylation efficiency was opposite between HL and LL plants. The decrease in photosynthetic efficiency in LL plants under low [O_2_] supports a more important role of O_2_ in photoprotection of LL plants (Sagun *et al.*, 2021). A link between electron transport and *R*_LIGHT_ is known to exist (Buckley and Adams, 2011), but it is not characterised in C_4_ plants. Of the fitted quantities, only *R*_LIGHT_ and *g*_BS_ were retained for model parameterisation, detailed above.

### Modelling of light distribution, light reactions and dark metabolism

Using measured anatomical characteristics, BS/M chlorophyll ratio and leaf absorptance (Table 1), we modelled light penetration through the leaf profile according to Bellasio and BS Lundgren (2016). Absorbed light in the BS relative to M was higher in HL plants (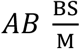 was 0.414 for HL and 0.343 for LL, Table 1). Next, to gain comprehensive understanding of the photosynthetic metabolism, we used these values, *Y*(*II*), *g*_BS_ and *R*_LIGHT_ derived from the concurrent gas-exchange and chlorophyll fluorescence measurements (Table S1), to parameterise a purposely derived model of light reactions and dark metabolism, while the fraction of electron flow through PSI following CEF in M and BS *f*_Cyc M_ and *f*_Cyc BS_, respectively) were fitted to maximise assimilation, constrained to maintain PPDK and PEPCK activity in BS cells at zero to match immunoblotting (Figure 5).

**Figure 5.**
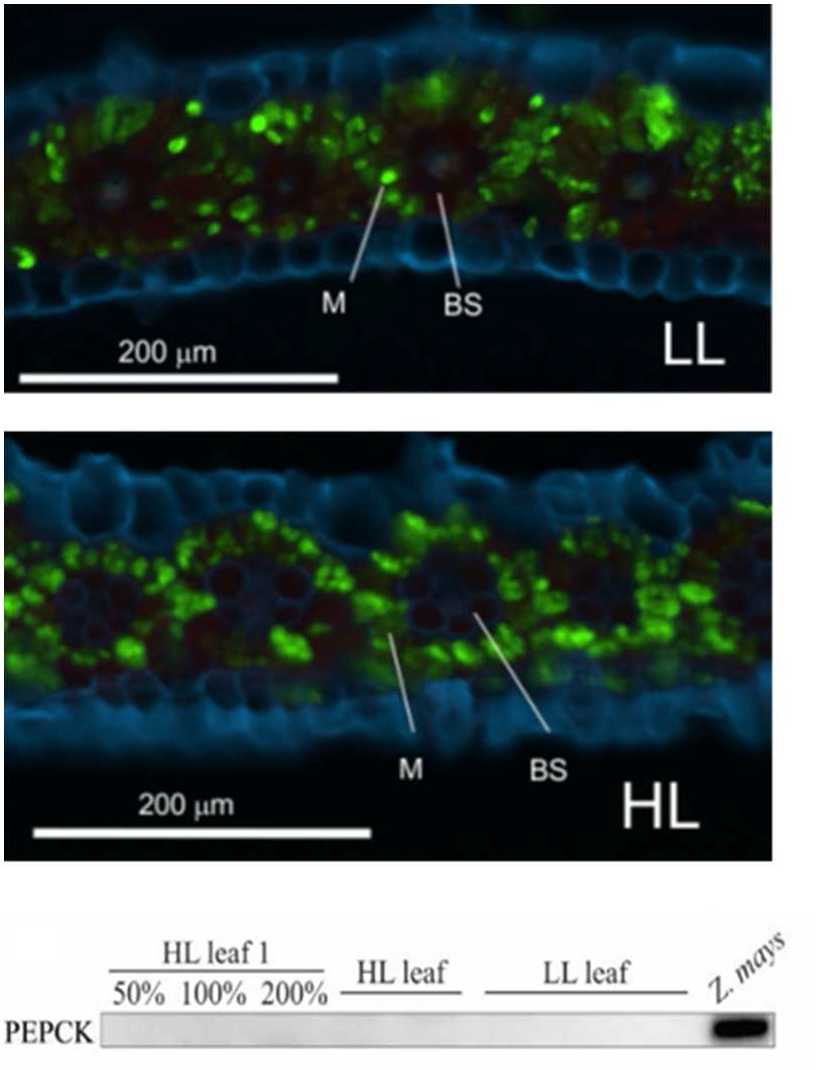
Protein assays. Fluorescent micrographs of pyruvate ortophosphate dikinase (PPDK) localisation on the leaf cross-sections of *S. viridis* grown at low light (LL) and high light (HL). Fluorescence signals are pseudo-coloured: green – PPDK labelled with secondary antibodies conjugated with Alexa Fluor 488; blue – calcofluor white-stained cell walls. BS, bundle sheath cells; M, mesophyll cells. At the bottom, immunodetection of PEP carboxykinase (PEPCK) in leaf protein extracts of *S. viridis* loaded on leaf area basis; *Z. mays* leaf sample was used for positive control. Three biological replicates were loaded for HL and LL plants.

Initially, we replicated *in silico* the leaf gas–exchange measurements under ambient O_2_ level (Figure 3). For both light and CO_2_ response curves, the trends of CO_2_ assimilation and *Y*(*II*) were well captured by the model. For assimilation, R^2^ was 0.98 and 0.97 in light curves or 0.95 and 0.81 in *A/C*_i_ curves of HL and LL plants, respectively. For *Y*(*II*), R^2^ was 0.86 and 0.96 in light curves or 0.91 and 0.76 in *A*/*C*_i_ curves of HL and LL plants, respectively. In the light response curves, the model captured a hyperbolic decrease of the measured *Y*(*I*) at high irradiance (Figure 3e) but not an initial dip at low irradiance. However, the modelled *Y*(*I*) was similar to the trend of 1 - *Y*(*ND*), perhaps only capturing *Y*(*I*) available due to the electron input from PSII but not accounting for the loss of *Y*(*I*) due to the unavailability of acceptors, that is, *Y*(*NA*). In CO_2_ response curves, the hyperbolic increase of *Y*(*I*) was captured, but *Y*(*I*) was slightly underestimated for *C*_M_ greater than 20 μmol mol^-1^. Predicted *Y*(*I*) did not differ between HL and LL plants.

Next, we carried out a simulation of photosynthetic biochemistry using the specific parameterisation of growth light regimes (Table 2). LL plants had a 60 % downregulation of assimilatory metabolism due to the lower energy availability (Table 3, to aid comparison between light regimes we also expressed fluxes in multiples of gross assimilation). Because of the lower NADPH availability, LL plants exported a higher fraction of the total PGA produced to the M for reduction than HL plants (42% and 35%, respectively, Table 3). Rubisco carboxylation (*V*_C_) and PEP carboxylation rate (*V*_P_, which corresponds to the rate of malate export to BS cells and the backflux of pyruvate) had lower activity in LL plants. The decrease in *V*_P_ caused a 25 % decrease in CO_2_ concentration in the BS that was reflected by a higher ratio of Rubisco oxygenation relative to carboxylation (*V*_O_/*V*_C_, 4.9 % in HL plants and 5.4 % in LL plants, Table 3). However, there was a general increase in PSII efficiency in LL plants which overweighed the effect on *V*O/*V*C. As a result, in LL plants the average quantum yield increased 38 % on absorbed light basis (from 0.04 to 0.055 CO_2_/quanta), and as much as 43 % on incident light basis (from 0.031 to 0.044 CO_2_/quanta) as compared to HL plants. LL plants gained more on incident light basis because of their higher leaf absorbance (captured in the measurements by *s*′, Table S1, and in the model inputs by *s*_0_, Table 2).

**Table 2.**
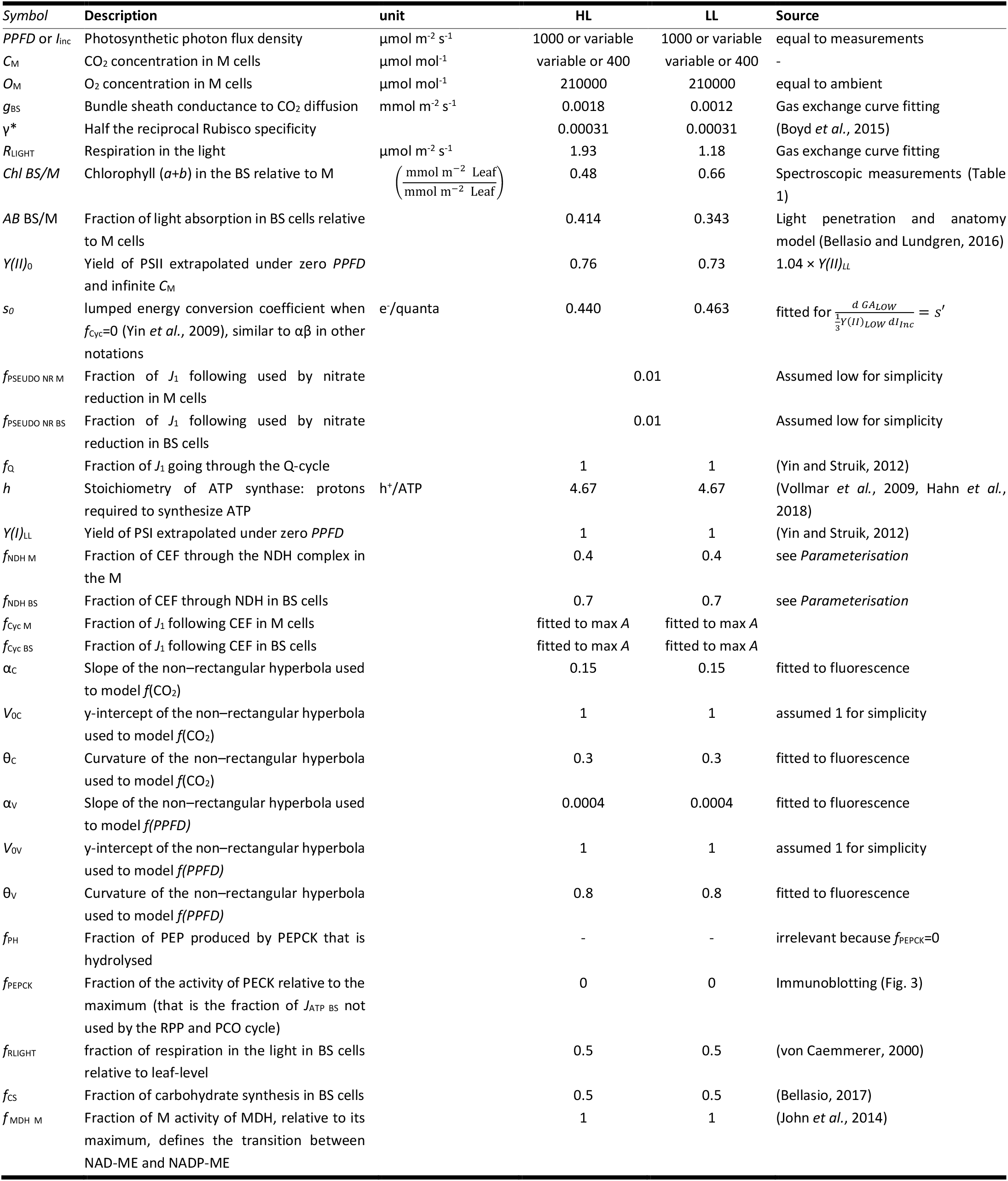
Model inputs and sources.

**Table 3.** Output of the combined light reaction and biochemical model of C_4_ photosynthesis. The output shown is calculated at the growth *PPFD* (1000 for HL or 300 μmol m^-2^ s^-1^ for LL), and expressed, except for *f*_Cyc_ and Φ (dimensionless) or *C* (μmol mol^-1^), in μmol m^-2^ s^-1^ and in brackets as a fraction of gross assimilation at the growth light (30.97 for HL and 13.24 μmol m^-2^ s^-1^ for LL plants).

| Symbol | Description | HL |  | LL |  |
| --- | --- | --- | --- | --- | --- |
|  |  | M | BS | M | BS |
| $f_{\text{Cyc}}$ | Fraction of electron flow through PSI ($J_1$) following CEF | 0.01 | 0.783 | 0.08 | 0.866 |
| $C$ | CO<sub>2</sub> concentration | 400 | 4100 | 400 | 3100 |
| $I_{\text{Inc}}$ | Incident light (*PPFD*) | 707 (23.5) | 293 (9.71) | 223 (17.7) | 77 (6.08) |
| $I_2$ | Light absorbed by PSII | 310 (10.0) | 50 (1.63) | 100 (7.54) | 9.5 (0.718) |
| $I_1$ | Light absorbed by PSI | 238 (7.68) | 176 (5.69) | 79.1 (5.97) | 51.9 (3.92) |
| $J_2$ | Electron flow through PSII | 151 (4.88) | 25 (0.794) | 63.8 (4.82) | 6.07 (0.458) |
| $J_1$ | Electron flow through PSI | 153(4.92) | 113 (3.65) | 69.2 (5.22) | 45.4 (3.43) |
| $J_{\text{NADPH}}$ | NADPH production rate (half the electron flow to NADPH) | 58.8 (1.90) | 7.09 (0.229) | 27.4 (2.07) | 0.948 (0.07) |
| $J_{\text{ATP}}$ | ATP production rate | 97.9 (3.16) | 80.2 (2.59) | 44.2 (3.34) | 32.5 (2.46) |
| $V_p$ | PEP carboxylation rate. Because in this study $f_{\text{MDHM}}=1$, $V_p$ corresponds to the rate of malate flux to the BS, the rate of pyruvate flux to the M, the PPK reaction rate in the M (PPDK reaction rate in the BS is fitted to be zero in this study) and the malic enzyme reaction rate | 36.7 (1.18) | 0 | 15.9 (1.20) | 0 |
| $L$ | Leakage rate of CO<sub>2</sub> out of the BS | 6.64 (0.214) | | 3.21 (0.242) | |
| $\Phi$ | Leakiness: CO<sub>2</sub> leakage rate out of the BS relative to $V_p$ | 0.18 | | 0.20 | |
| $V_c$ | Rubisco rate of carboxylation | 0 | 31.7 (1.02) | 0 | 13.6 (1.03) |
| $V_o$ | Rubisco rate of oxygenation | 0 | 1.54 (0.0496) | 0 | 0.735 (0.0555) |
| $PR$ | Rate of PGA reduction | 22.2 (0.716) | 43.0 (1.39) | 11.5 (0.868) | 16.4 (1.24) |
| $DHAP_{\text{RPP}}$ | Rate of DHAP entering the conversion phase of the RPP cycle | 0 | 55.5 (1.79) | 0 | 23.9 (1.81) |
| $RuP_{\text{Phosph}}$ | Rate of RuP phosphorylation | 0 | 33.3 (1.07) | 0 | 14.3 (1.08) |
| $CS$ | Rate of carbohydrate synthesis | 4.85 (0.156) | 4.85 (0.156) | 2.01 (0.152) | 2.01 (0.152) |
| $R_{\text{Light}}$ | Rate of respiration in the light | 0.95 (0.031) | 0.95 (0.031) | 0.590 (0.047) | 0.590 (0.046) |
| PGA flux | Flux of PGA | 22.5 (0.726) | -22.5 (-0.726) | 11.7 (0.882) | -11.7 (-0.882) |
| DHAP flux | Flux of DHAP | -17.3 (-0.559) | 17.3 (0.559) | -9.48 (-0.716) | 9.48 (0.716) |

## Discussion

*S. viridis* is a wild ancestor of *S. italica* (foxtail millet), a grain crop widely grown in China and India. It has gained favour for studying C_4_ photosynthesis because it has a rapid life cycle, small stature, sequenced genome and available genetic transformation method (Brutnell *et al.*, 2010, Ermakova *et al.*, 2021c). Since C_4_ photosynthesis has evolved independently at least 70 times (Sage, 2017), diverse biochemical arrangements are possible in C_4_ plants, and characterising different approaches of shade acclimation is a prerequisite for identifying best targets for improving crop performance. While anatomical responses of C_4_ plants to shading are relatively conserved (Ward and Woolhouse, 1986, Pengelly *et al.*, 2010), biochemical strategies appear to be diverse.

We recently showed that *S. viridis* grown at limiting irradiance deployed a suite of protein level adjustments providing BS thylakoids with increased capacity for light harvesting and electron transport and pointing to an increased ATP demand in the BS (Ermakova *et al.*, 2021b). Here, we were interested in resolving whether the observed rearrangements were triggered by an increased ATP consumption from dark reactions or by a lower rate of production, perhaps, due to anatomical modifications. Whilst acclimation to LL generally entails a downregulation of photosynthetic potential, here gas exchange measurements showed that HL and LL plants reached strikingly similar rates of assimilation when measured at the same light or CO_2_ levels. This intriguing invariance, which we repeatedly measured in plants from different growth batches, and was previously observed in the same ecotype independently (Henry *et al.*, 2019), may be a particular feature of *S. viridis.* Detailed analysis of the response curves through model fitting (Table S1) revealed some contrasting acclimation strategies: LL plants had lower carboxylation efficiency and lower *Y*(*II*)_LL_, but also lower respiration, than HL plants. We then characterised the leaf anatomy and found a reduction in BS size and in BS surface area per leaf area (Sb) in LL plants. Our interpretation is that these anatomical changes concurred to the reduction of *g*_BS_ (von Caemmerer *et al.*, 2008), here estimated by fitting of gas exchange and fluorescence data obtained under ambient and low [O_2_]. The reduction of *g*_BS_ at low light is consistent with previous results obtained through different proxies including anatomy, isotopic discrimination, combined gas exchange and chlorophyll fluorescence (Pengelly *et al.*, 2010, Ubierna *et al.*, 2013). Interestingly, *g*_BS_ was shown to decrease both in plants grown under low light (Bellasio and Griffiths, 2014b) and in plants grown under high light subsequently transferred to low light, as it occurs in crop canopies where older leaves are shaded by new growth (Bellasio and Griffiths, 2014a).

The decrease in *g*_BS_ ameliorates the efficiency of C_4_ photosynthesis. By hindering CO_2_ leakage, it counters the decrease in CO_2_ concentration in the BS and the increase of photorespiration occurring under low light (Kromdijk *et al.*, 2014). However, the associated reduction in BS size (Table 1) has the undesirable consequence of decreasing the light harvesting in BS cells, limiting ATP generation and decreasing the operational plasticity under changing light conditions (Bellasio and Lundgren, 2016). LL plants deployed a concerted suite of responses to counter potential ATP starvation in the BS. Firstly, while in HL plants M chloroplasts were dispersed, in LL plants the arrangement of M chloroplasts along the cell walls (Figure 1), similar to that observed in *Saccharum officinarum* grown under low light (Sales *et al.*, 2018), created optical gaps, resulting in lower apparent absorbance of the M and increasing the amount of light filtering through to BS cells (Table 1), a phenomenon known as ‘sieve effect’ (Terashima *et al.*, 2009). Chloroplast arrangements differ depending on species, light quality and intensity. For instance, under midday illumination *Eleusine coracana* was shown to aggregate M chloroplasts around the BS; *Zea mays* showed a dispersed arrangement, similar to what we observed in *S. viridis*; *Sorghum bicolor* created characteristic optical corridors spanning the whole leaf thickness, presumably aiding photoprotection, observed also in *Z. mays* but only under non–physiological irradiance (3,000–4,000 μmol m^-2^ s^-1^) (Yamada *et al.*, 2009, Maai *et al.*, 2020a, Maai *et al.*, 2020b). Secondly, in LL leaves the BS had higher chlorophyll content (Table 1) and increased abundance of light-harvesting complexes (LHC) I and II compared to HL plants (Ermakova *et al.*, 2021b). Consistently, LL leaves had higher absorptance (Table 1), and fitting of combined gas exchange and fluorescence data captured a higher *s*′, the parameter representing effective leaf level light interception lumped to the yield of ATP generation (Table S1). Thirdly, the potential of BS thylakoid membranes for ATP production increased due to the upregulation of PSI, Cytochrome *b*_6_*f* ATP synthase and NDH (Ermakova *et al.*, 2021b). The latter has been shown to be beneficial for C_4_ photosynthesis in reverse genetics studies (Ishikawa *et al.*, 2016, Peterson *et al.*, 2016) and found to correlate with the ATP requirements of cell types in C_4_ plants (Takabayashi *et al.*, 2005).

Next, we were interested in resolving to what extent the observed reorganisations of light harvesting and electron transport processes in LL plants was effective in countering the limitations to ATP generation caused by a smaller BS, and what modifications to the metabolism of BS and M were required to maintain invariant levels of assimilation (Figure 3). We developed a model encompassing explicit anatomy and separate electron transport chains in the BS and M, parameterised specifically for HL and LL plants. We found that the increased chlorophyll concentration was not sufficient to counter the effect of smaller BS cells size and, overall, LL plants absorbed 40 % less light in BS cells relative to total (from 23 % under HL to 14 % under LL, Table 3). However, the higher proportion of CEF *f*_Cyc_) in BS of LL plants (Table 3, Figure S3) could partially compensate for the decrease in light absorption, so that the predicted ATP production rate, relative to total, only decreased by about 5 % (from 45 % in the BS of HL plants to 42 % in the BS of LL plants, Table 3). The predicted changes in *f*_Cyc_ – an emerging property of the fitted model mainly resulting from the reduction in BS size – are in line with the increased content of NDH detected in LL BS cells (Ermakova *et al.*, 2021b), again pointing to the key role of NDH in ATP production in BS cells. Hence, we demonstrated that: first, the upregulation of CEF *f*_Cyc_) is effective in mitigating decreased light interception in BS cells, and, second, that decreasing light interception in BS cells is a sufficient condition to require an increase of *f*_Cyc_ (Table 3, Figure S3). To our knowledge this functional dependence was not demonstrated before, uncovering another important link between biochemical properties and structural characteristics of C_4_ leaves.

The NADPH production rate in BS cells relative to total was predicted to decrease 97 % in LL plants (from 11 % under HL to 0.3 % under LL), driven both by the decrease in BS size and increase in *f*_Cyc_. This led to a 13 % decrease in the rate of PGA reduction in BS cells relative to total (from 66 % under HL to 59 % under LL), and a consequent relative increase of triose fluxes between M and BS cells in LL plants (Table 3). By means of LEF, PSII activity in BS cells not only supplies NADPH production, but also replenishes CEF electrons leaking to various electron sinks downstream of PSI (explicitly accounted for in the model as pseudocyclic electron flow, but not varied in this study). In this model output, LEF in the BS was predicted to be 21 μmol m^-2^ s^-1^ (14 % of the total leaf rate for HL plants), and 5.59 μmol m^-2^ s^-1^ (9 % of the total leaf rate for LL plants, Table 3). The actual PSII activity in the BS thylakoid membranes measured by Ermakova *et al.* (2021b) matched the value predicted for LL plants (it was circa 7 % of the leaf rate), but in HL plants it was a half of the LL value, much lower than the model prediction. This suggests that the predicted value of *J*_2_ in the BS of HL plants cannot be supplied *in vivo* by electrons coming from water oxidation, but must be obtained via oxidation of NADPH, derived from malate imported from the M, through the NDH. This idea is supported by experimental observations. When supplied with malate, isolated BS strands of *Z. mays* retained PSI activity even in the presence of PSII inhibitor. Further, the BS of *S. bicolor* maintained CO_2_ fixation under far–red illumination, which is unable to excite PSII (Osmond, 1974, Ivanov *et al.*, 2005).

Theory shows that, under changing irradiance, flexibility in the apportioning of PGA reduction between BS and M cells can regulate the BS demand of NADPH, and the engagement of PEPCK and PPDK can attune the BS demand of ATP (Furbank, 2011, Bellasio and Griffiths, 2014c). Because PEPCK consumes ATP generated in the M and is tightly regulated by ATP availability, the expression of PEPCK in NADP–ME plants was predicted to increase light harvesting plasticity, *i.e.*, the capacity to efficiently harvest light in a broad range of intensity and spectral quality (Bellasio and Griffiths, 2014c). Further, the activity of PPDK in the BS could compensate for a lack of PEPCK, making PEPCK engagement necessary when the activity of PPDK in the BS was reduced, and *vice versa* (Bellasio, 2017). Indirect evidence supports these predictions. It was shown that low light acclimated NADP–ME plants, like *Z. mays* and *S. officinarum*, increased the total rate of PEPCK activity (Sales *et al.*, 2018) or the activity of PEPCK relative to PEPC (Sharwood *et al.*, 2014, Sonawane *et al.*, 2018), and increased the pool of inactive PEPCK (data made available by courtesy of B.V. Sonawane, personal communication), presumably available to be activated by changing light conditions (Bailey *et al.*, 2007).

Uniquely, *S. viridis* did not express PEPCK neither under HL nor under LL (Figure 5). Nor we found detectable levels of PPDK in the BS (Figure 5), in line with previous reports (John *et al.*, 2014, Schlüter and Weber, 2020). Although *Z. mays* grown under low light overexpressed LHCII subunits and PsbD in BS cells (Drozak and Romanowska, 2006), it did not show increased oxygen activity or changes in supramolecular organisation of PSII (Romanowska *et al.*, 2006, Rogowski *et al.*, 2019). In contrast, *S. viridis* had a striking plasticity in acclimating light reactions, probably sufficient to avoid the necessity to regulate PEPCK and PPDK expression (Figure 5). Importantly, and differently from the nine species (four NADP–ME including *Z. mays* and *S. bicolor*, two NAD–ME, two PEPCK) studied by Sonawane *et al.* (2018), acclimation of light reactions in *S. viridis* did not compromise quantum yield (Table S1). Therefore, the outstanding capacity of *S. viridis* to rearrange light reactions under low light is a highly efficient acclimation strategy setting *S. viridis* aside from other NADP–ME plants and representing a so far overlooked innovation in C_4_ evolutionary history.

## Conclusion

Using a purposely developed model, parametrised with original anatomical, biochemical, and gas exchange data, we analytically and mechanistically solved the causality link between anatomy and biochemistry. The optical cross section determined the absorption of light in the BS and the M, the rate of electron transport, the engagement of CEF and LEF, the rate of ATP and NADPH generation, and ultimately the apportioning of carbon metabolism between M and BS cells. To counter ATP starvation, *S. viridis* adjusted electron transport processes, boosting cycling electron flow in the BS. Although quantifying the impact in field conditions will require additional experiments, the striking capacity of *S. viridis* to acclimate light reactions is perhaps the most efficient shade tolerance strategy, potentially leading to novel possibilities for crop improvement.

## Availability

Data and the model, coded in **R** and Microsoft® Excel®, are made freely available in the Supporting Information and on GitHub at the link [to be provided during copyediting].

## Author contributions

ME and CB conceived the project. CB developed the model and ran simulation. CB and ME performed experiments, analysed the data and wrote the article. Authors declare no conflict of interest.

## Acknowledgements

We are deeply grateful to Graham Douglas Farquhar for hospitality and to Suan Chin Wong, Dean Price and Susanne von Caemmerer for equipment. We thank Russell Woodford for coding in R, Balasaheb Vitthal Sonawane for data and discussion, Riya Kuruvilla and Tegan Norley for technical assistance. We thank Florence Danila, Joanne Lee and Centre for Advanced Microscopy at the Australian National University for preparing and imaging leaf cross-sections. We thank the Australian Plant Phenomics Facility supported under the National Collaborative Research Infrastructure Strategy of the Australian Government. CB was funded by H2020 Marie Skłodowska-Curie individual fellowship (DILIPHO, ID: 702755). ME was supported by the Australian Research Council Centre of Excellence for Translational Photosynthesis (CE140100015).

